# The Role of the Wnt/PCP Formin Daam1 in Renal Ciliogenesis

**DOI:** 10.1101/512533

**Authors:** Mark E. Corkins, Vanja Krneta-Stankic, Malgorzata Kloc, Pierre D. McCrea, Andrew B. Gladden, Rachel K. Miller

## Abstract

Kidneys are composed of numerous ciliated epithelial tubules called nephrons. Each nephron functions to reabsorb nutrients and concentrate waste products into urine. Defects in primary cilia are associated with abnormal formation of nephrons and cyst formation in a wide range of kidney disorders. Previous work in *Xenopus laevis* and zebrafish embryos established that loss of components that make up the Wnt/PCP pathway, Daam1 and ArhGEF19 (wGEF) perturb kidney tubulogenesis. Dishevelled, which activates both the canonical and non-canonical Wnt/PCP pathway, affect cilia formation in multiciliated cells. In this study, we investigated the role of the noncanoncial Wnt/PCP components Daam1 and ArhGEF19 (wGEF) in renal ciliogenesis utilizing polarized mammalian kidney epithelia cells (MDCKII and IMCD3) and *Xenopus laevis* embryonic kidney. We demonstrate that knockdown of Daam1 and ArhGEF19 in MDCKII and IMCD3 cells leads to loss of cilia, and Daam1’s effect on ciliogenesis is mediated by the formin-activity of Daam1. Moreover, Daam1 co-localizes with the ciliary transport protein IFT88. Interestingly, knocking down Daam1 in *Xenopus* kidney does not lead to loss of cilia. This data suggests a new role for Daam1 in the formation of primary cilia.

## INTRODUCTION

Primary cilia are microtubule-based cellular protrusions that allow a cell to sense its environment (Satir *et al.*, 2010). Many cell types in the body contain cilia, and improper cilia development results in a family of diseases called ciliopathies, including polycystic kidney disease, nephronophthisis, Joubert syndrome and Bardet-Biedel syndrome (Lee and Gleeson, 2011). Although ciliopathies can manifest in a number of different ways, the vast majority of ciliopathies result in kidney abnormalities (Arts and Knoers, 2013). For example, one of the first described ciliopathies is the result of a mutation in the gene IFT88*/*Polaris, a protein is transported within vesicles to facilitate ciliary biogenesis (Yoder *et al.*, 2002; Ding *et al.*, 2017). Loss of IFT88 leads to cystic kidney disease in both mouse and human (Moyer *et al.*, 1994; Onuchic *et al.*, 1995). Additionally, loss of IFT88 in human mesenchymal stem cells leads to increased Wnt signaling (McMurray *et al.*, 2013).

The canonical Wnt pathway has many connections to cilia development. In the zebrafish left/right organizer, loss of β-catenin, the Wnt co-transcription factor, results in a reduction of cilia (Zhu *et al.*, 2015). In mouse and *Xenopus laevis* a protein called Chibby1, which functions to shuttle β-catenin out of the nucleus, localizes to the cilia and is required for proper ciliogenesis and kidney development (Lee *et al.*, 2014; Shi *et al.*, 2014). Additionally, mutations in genes that regulate cilia formation typically results in increased sensitivity to Wnt ligands, although the mechanism for this phenomenon is heavily debated (Corbit *et al.*, 2008). Moreover, Dishevelled, a component of both the canonical and non-canonical Wnt pathway, is required for actin assembly and positioning of the basal bodies on the surface of the cell during ciliogenesis in *X. laevis* multiciliated skin cells (Park *et al.*, 2008).

Actin filaments are important for proper ciliogenesis. In motile multiciliated cells, the cilia are connected by an actin network (Werner *et al.*, 2011). Although this actin network is not clearly visible around primary cilia, application of drugs that broadly inhibit or stabilize actin filaments typically results in longer primary cilia (Drummond *et al.*, 2018). Formin proteins act to polymerize actin filaments. However, treatment with the formin inhibitor smiFH2 has been found to decrease the number and length of primary cilia (Copeland *et al.*, 2018). This suggests a complex story in which different sub-populations of actin can have either a positive or negative affect on ciliogenesis.

Daam1 (Dishevelled Associated Activator of Morphogenesis 1) is a Diaphanous-related Formin homology (DRF) protein that functions through the non-canonical Wnt/PCP pathway. It is regulated in part by Wnt ligands through Frizzled receptors and Dishevelled (Liu *et al.*, 2008). Daam1 is associated with a number of cellular functions, such as directed cell migration, planar cell polarity (PCP) and endocytosis (Kida *et al.*, 2007; Hoffmann *et al.*, 2014; Luo *et al.*, 2016).

Developmentally, Daam1 is necessary for gastrulation and normal kidney development in *X. laevis* (Habas *et al.*, 2001; Miller *et al.*, 2011; Corkins *et al.*, 2018). Additionally, it is present in the multiciliated epidermal cells of the *X. laevis* skin where it regulates the actin network that stabilizes cilia. Loss of Daam1 in motile cilia results in their loss of polarity (Yasunaga *et al.*, 2015).

The Diaphanous family of formin proteins is regulated by the Rho family of GTPases (Alberts, 2001). Daam1 is known to bind to RhoA, a protein that regulates cytoskeletal dynamics. However unlike other Diaphanous family formin proteins, Daam1 is thought to be more strongly regulated by Dishevelled than Rho (Liu *et al.*, 2008). Wnt signaling through Dishevelled can activate RhoA, a process that is regulated by the Daam1/ArhGEF19 (wGEF) complex (Tanegashima *et al.*, 2008). In mice, *arhgef19* is expressed mainly in the intestine, liver, heart and kidney (Wang *et al.*, 2004), and loss of either ArhGEF19 or Daam1 in *X. laevis* leads to kidney malformations (Miller *et al.*, 2011; Corkins *et al.*, 2018).

In this study, we find that loss of the PCP component Daam1 negatively regulates ciliogenesis in MDCKII and IMCD3 cells. Daam1 knockdown within these mammalian kidney epithelial cells result in the formation of fewer cilia. Furthermore, knockdown of Daam1 within *Xenopus* embryonic kidneys does not appear to result in disruption of primary ciliogenesis. Daam1 deficient kidneys have primary cilia; however, we cannot exclude the possibility that their structure, organization and/or function is not affected. Our data indicate that ciliogenesis in MDCKII cells depends on the formin activity of Daam1. Additionally, the Daam1 partner, ArhGEF19, is also required for proper cilia formation in MDCKII cells. Finally, we found that in MDCKII and IMCD3 cells Daam1 localizes in vesicles that carry ciliary components, further supporting its role in ciliogenesis.

## RESULTS

### Loss of Daam1 results in a failure of kidney epithelial cells to ciliate

To determine whether Daam1 is required for primary ciliogenesis, *daam1* was knocked down in polarized MDCKII cells. MDCKII kidney epithelia form primary cilia when plated on a transwell membrane and are allowed to grow until confluency. Loss of Daam1 leads to cilia reduction in MDCKII cells (Figure 1). To ensure that this phenotype is due to loss of Daam1 we used two different shRNAs: one targeting the 3’UTR (#1) and one targeting the coding sequence of *daam1* (#3). Both shRNAs reduce Daam1 protein expression as visualized by Western blot (Figure 1E). A third shRNA, *sh-daam1* #2, failed to reduce Daam1 expression (Figure 2C). Additionally, similar experiments were carried out in IMCD3 cells using a *sh-daam1* #1 construct adapted for mice. These experiments also indicate that knockdown of *daam1* disrupts primary ciliogenesis (Figure S1A,B).

**Figure 1:**
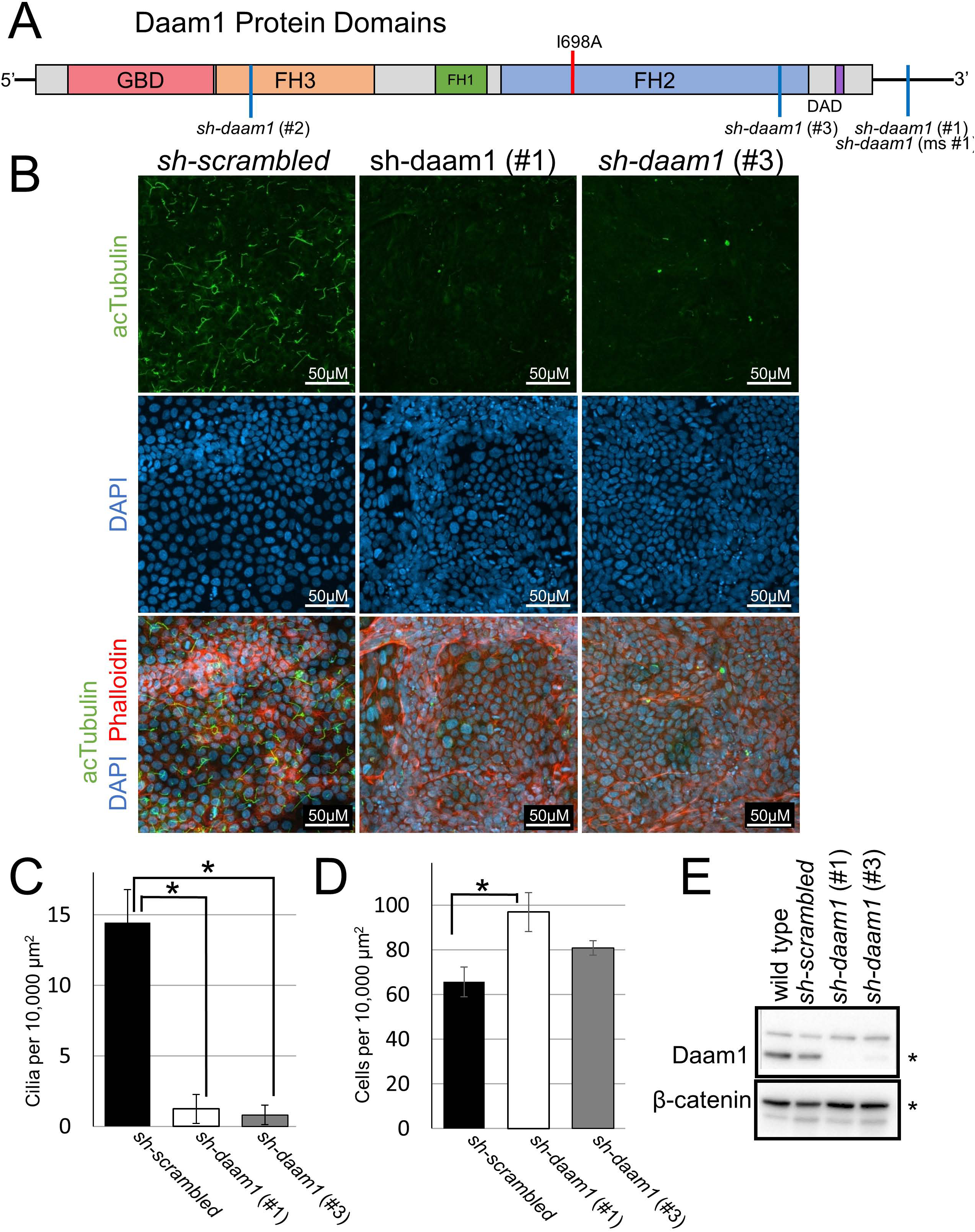
Daam1 knockdown results in a loss of primary cilia in MDCKII cells. **A**) Diagram illustrating the domains within the Daam1 protein. The GBD(DiaphanousGTPase-binding Domain) FH3(Diaphanous FH3 Domain) FH1(Formin Homology 1) FH2(Formin Homology 2) and DAD(Diaphanous Auto-regulatory Domain) are indicated. The location of the I698A mutation within the formin homology 2 domain (FH2) is marked with a red line, and positions corresponding to the shRNAs are marked with a blue line. MDCKII canine kidney epithelial cells were infected with either *sh-daam1* or a control construct and grown upon transwell filters. **B)** Cells were stained with acetylated α-Tubulin antibody (acTubulin) to visualize primary cilia (green), DAPI to label nuclei (blue), and phalloidin to label F-actin (red). Confocal imaging was used to analyze the effect of Daam1 depletion on primary ciliogenesis. Scale bars equal to 50 μm. **C)** Cilia were quantified. Error bars are shown as ± SEM. **D)** Cell numbers were quantified as described in Figure S5. Error bars are shown as ± SEM. **E)** Western blot of *sh-daam1* MDCKII cell lysates showing depletion of Daam1 protein levels. β-catenin was used as a loading control. * indicates p<0.05 as compared to *sh-scrambled*.

**Figure 2:**
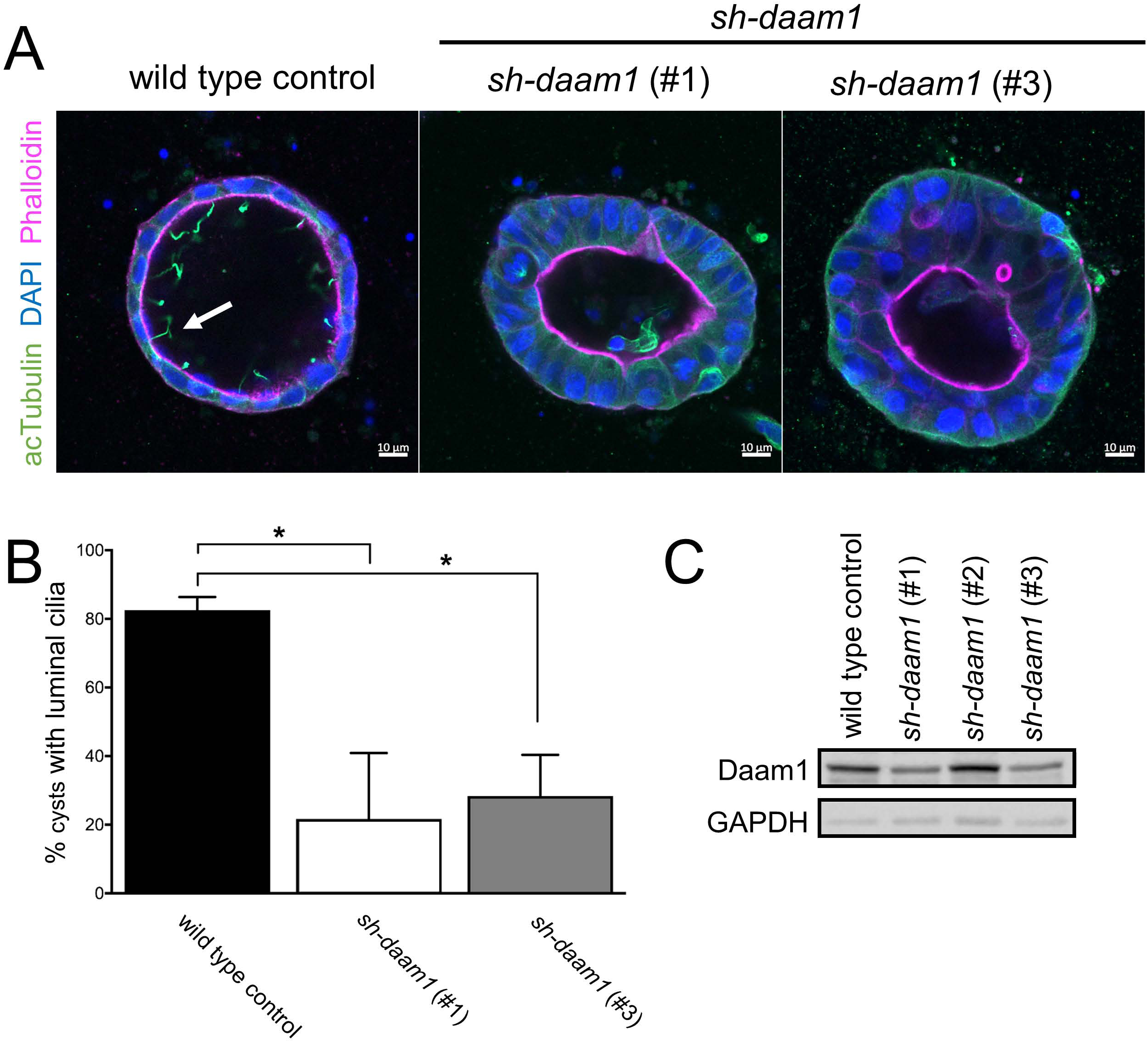
*sh-daam1* depleted MDCKII cells show reduced luminal ciliogenesis in 3D cultures. Control and *sh-daam1*-depleted cells were cultured in collagen I matrix to form cysts. After 12-14 days in culture, cysts were fixed and stained with an antibody against acetylated α-Tubulin (acTubulin) to visualize primary cilia (green), phalloidin for F-actin (magenta) and DAPI for nuclei (blue). Using confocal imaging we analyzed the effect of Daam1 depletion on ciliogenesis. (**A)** Representative merged confocal images showing reduction of luminal cilia upon Daam1 knockdown. White arrow points to luminal cilia in control cysts. Scale bars equal to 10 μm. **(B)** The graph shows quantification of cysts with luminal cilia for each condition. Twenty randomly chosen cyst per sample were analyzed in three independent experiments. Error bars are shown as ± SEM; Significance was calculated using unpaired, two-tailed t-test; ns indicates p>0.05, * indicates p<0.05, **p<0.01 **(C)** Western blot showing Daam1 protein levels in wild-type (control) cells and in cells expressing different shRNAs targeting Daam1. GAPDH was used as loading control.

Not only do *sh-daam1* cells have fewer numbers of cilia, but the cells grow to a higher density and exhibit a tendency to grow on top of each other, rather than forming a cell monolayer. When wild type cells pile on top of each other, they tend not to ciliate. Therefore, in our data analysis, we avoided scoring areas where cells failed to form a single monolayer.

MDCKII cells are used extensively to study renal tubulogenesis because they are capable of forming tubule-like structures when cultured in three-dimensional (3D) gels. These structures appear as single-layered, polarized, and hollow spherical cysts that closely reassemble the architecture of nephric tubules. Epithelial cells within cysts display apico-basal polarity and form luminal cilia similar to that of a nephron (Baek *et al.*, 2016). Ciliogenesis is a complex process that may involve the interaction of cells with the extracellular matrix (ECM) (Bryant *et al.*, 2010). Thus, to further investigate the role of Daam1 in ciliogenesis, we cultured control and *sh-daam1* depleted cells in collagen I matrix.

Control and Daam1 knockdown cells both formed 3D cysts; however, Daam1-depleted cysts showed perturbed primary ciliogenesis (Figure 2A and B). Daam1-depleted cysts showed a significant reduction of luminal cilia compared to controls (Figure 2A and B). Moreover, Daam1-depleted cysts displayed increased ciliogenesis on the basal - ECM facing side (Figure S2A and B). These results suggest that Daam1 is involved in regulating ciliogenesis and may be important for establishment and/or maintenance of apico-basal polarity. Additionally, we found that Daam1-depleted cells were more likely to form single lumens and present with cells within luminal space in comparison with control (Figure S2A and B).

### Daam1 formin activity is required for ciliogenesis

Given prior studies indicating that the actin cytoskeleton that stabilizes multicilia in *X. laevis* skin, we hypothesized that the formin activity of Daam1 is required for ciliogenesis. Rescue experiments were carried out to investigate *sh-daam1* cilia phenotype using constructs that express either GFP-Daam1 or a GFP-Daam1(I698A) mutant, which disrupts the actin binding activity of the FH2 domain(Moseley *et al.*, 2006; Lu *et al.*, 2007). Due to significant toxicity upon stable overexpression of Daam1 we utilized transient transfections to express wild type Daam1 or Daam1(I698A).

Approximately 30% of the cells had strong GFP expression. Wild type Daam1 partially rescued the reduced cilia phenotype (Figure 3) In contrast, the Daam1(I698A) mutant failed to rescue the cilia phenotype.

**Figure 3:**
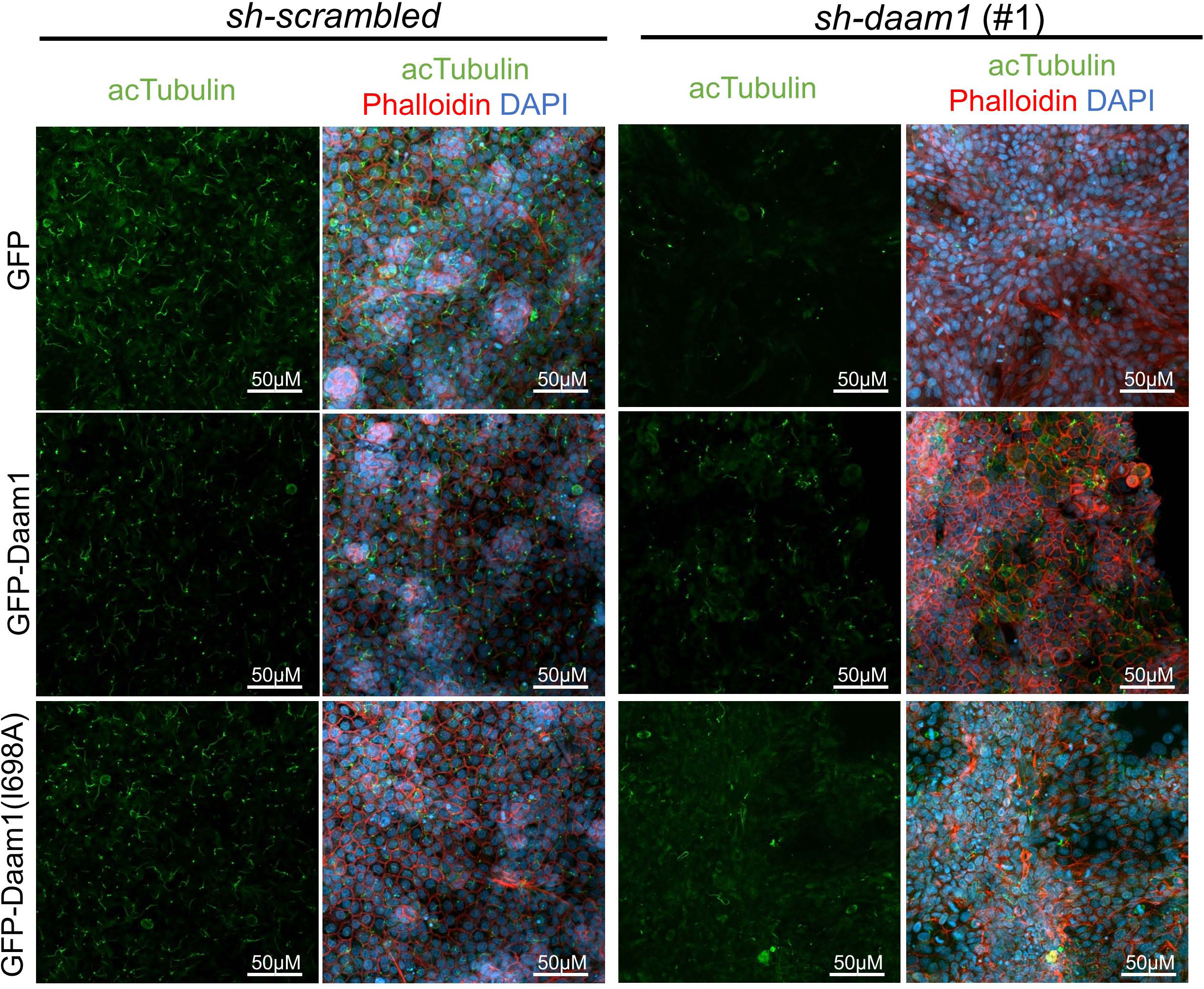
The *sh-daam1* cilia phenotype can be rescued with wild-type Daam1 but not Daam1(I698A). Stable MDCKII cells infected with either *sh-daam1* or a control *sh-scrambled* construct were transiently transfected with constructs that express either GFP, GFP-Daam1, or GFP-Daam1(I698A) constructs. Cells were grown on transwell filters and stained with acetylated α-Tubulin antibody (acTubulin) to label primary cilia (green), DAPI to label nuclei (blue), and phalloidin to label F-actin (red). Confocal imaging was used to analyze the effects upon primary ciliogenesis. Scale bars equal to 50 μm.

### Daam1 localizes in vesicles that carry ciliary components

To further examine the role of Daam1 in ciliogenesis, the subcellular localization of Daam1 was observed in live cells transfected with GFP-Daam1. Daam1 predominantly localizes to highly motile puncta (Figure 4, S1C and S3).

**Figure 4:**
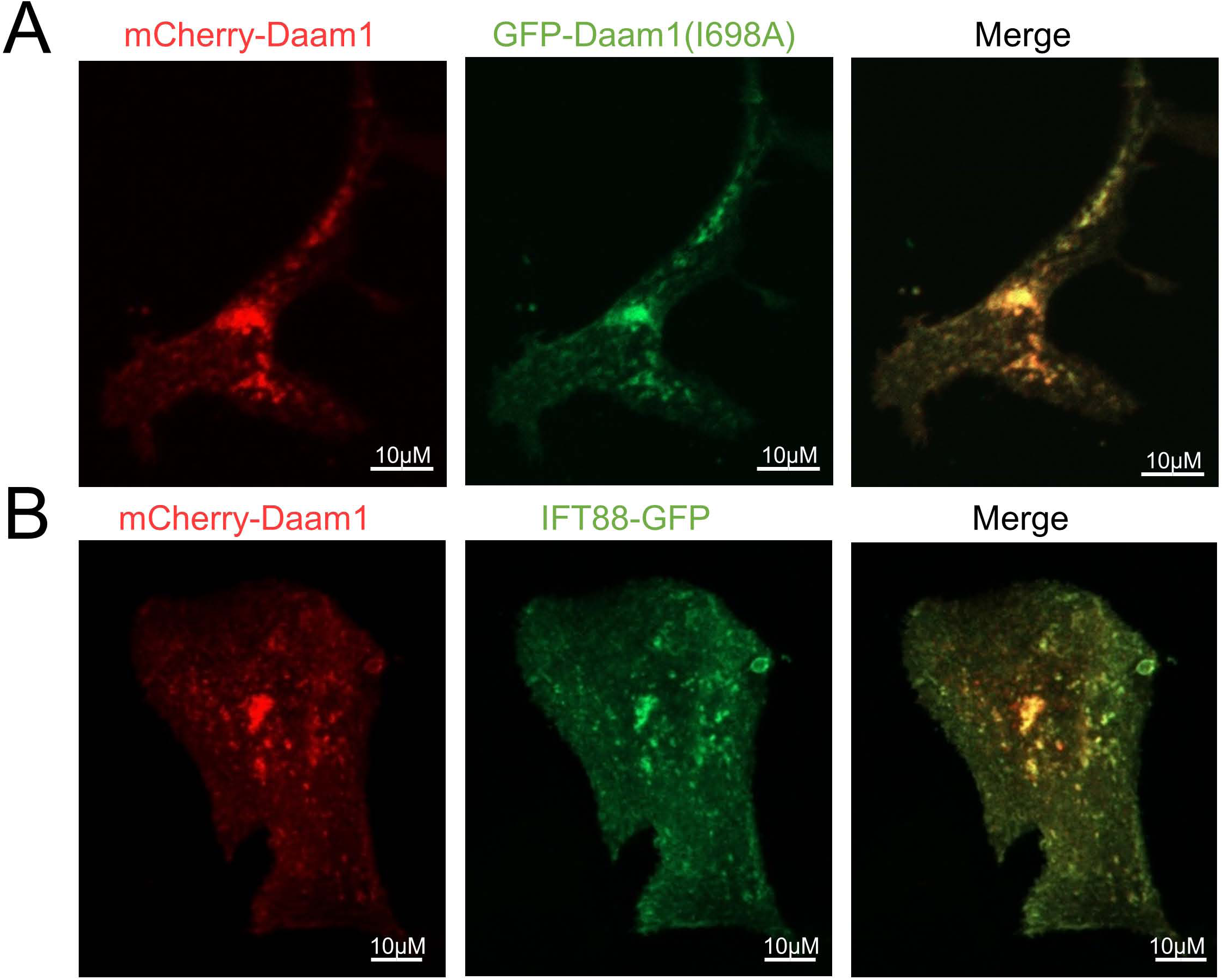
Daam1 localizes to vesicles that carry ciliary components. **A**) MDCKII cells were co-transfected with constructs that express mCherry-Daam1 and GFP-Daam1(I698A) then imaged via confocal to analyze the localization in live cells. **B)** MDCKII cells were co-transfected with constructs that express mCherry-Daam1 and IFT88-GFP than imaged via confocal in live cells to determine the subcellular localization. Scale bars equal to 10 μm.

Additionally, because the Daam1 formin mutant failed to rescue loss of cilia and some of the Daam1 truncations such as Daam1(524-1078) fail to localize properly (Luo *et al.*, 2016), GFP-Daam1(I698A) localization was visualized to determine whether it is similar to that of wild type Daam1. mCherry versions of these constructs were made, then co-transfected. Both the wild type and I698A versions were found to colocalize with each other (Figure 4A).

To assess whether Daam1 is localized at the cilia, its localization was visualized concurrently with that of ciliary markers. Given that fixation and staining resulted loss of GFP-Daam1 signal, live cells were utilized to assess Daam1 localization. However, because MDCKII cells are polarized on an opaque membrane and a high level of autofluorescence appears after 5 days of confluency, MDCKII cells are not ideal for live imaging. Therefore, transiently transfected IMCD3 were ciliated on glass coverslips over a shorter period of time to directly visualize cilia in live cells. A GFP-Chibby1 construct was used to label the transition zone of the cilia and an α-Tubulin-GFP construct was used to label the axoneme of the cilia (Figure S3) (Steere *et al.*, 2012). Using these constructs, co-localization of Daam1 with the ciliary transition zone or axoneme of cilia was not apparent.

Because vesicles are involved in cilium biogenesis, localization studies were carried out to determine whether Daam1 puncta are vesicles that carry ciliary components. A mCherry-Daam1 construct was co-transfected with an IFT88-GFP construct (Ding *et al.*, 2017). Daam1 localizes to vesicles that carry IFT88 (Figure 4B, S1C). Both the IFT88 and Daam1 labeled vesicles present in MDCKII cells are much smaller and more numerous than the vesicles in IMCD3 cells (Figures 4B, S1C). Daam1 is found in vesicles that are labeled with IFT88, a gene involved in ciliogenesis.

### ArhGEF19 is required for ciliogenesis

As loss of Daam1 results in loss of cilia, we assessed whether ArhGEF19, a RhoGEF that associates with Daam1 and acts within the non-canonical Wnt/PCP pathway, is required for ciliation. If RhoA activation is required for ciliogenesis, loss of ArhGEF19 should mimic the phenotypes shown in *sh-daam1* cells. Loss of *arhgef19* results in almost complete loss of cilia (Figure 5). Similar phenotypes were observed in *sh-arhgef19* cells and *sh-daam1* cells, with *sh-arhgef19* cells growing to higher density and exhibiting cilia loss. However, one distinction noted was that in contrast to *sh-daam1* cells, the *sh-arhgef19* cells do not grow on top of each other, indicating that distinct mechanism may also exist. The loss of cilia upon *arhgef19* depletion indicates that GTP-RhoA as part of the PCP pathway is necessary for ciliogenesis.

**Figure 5:**
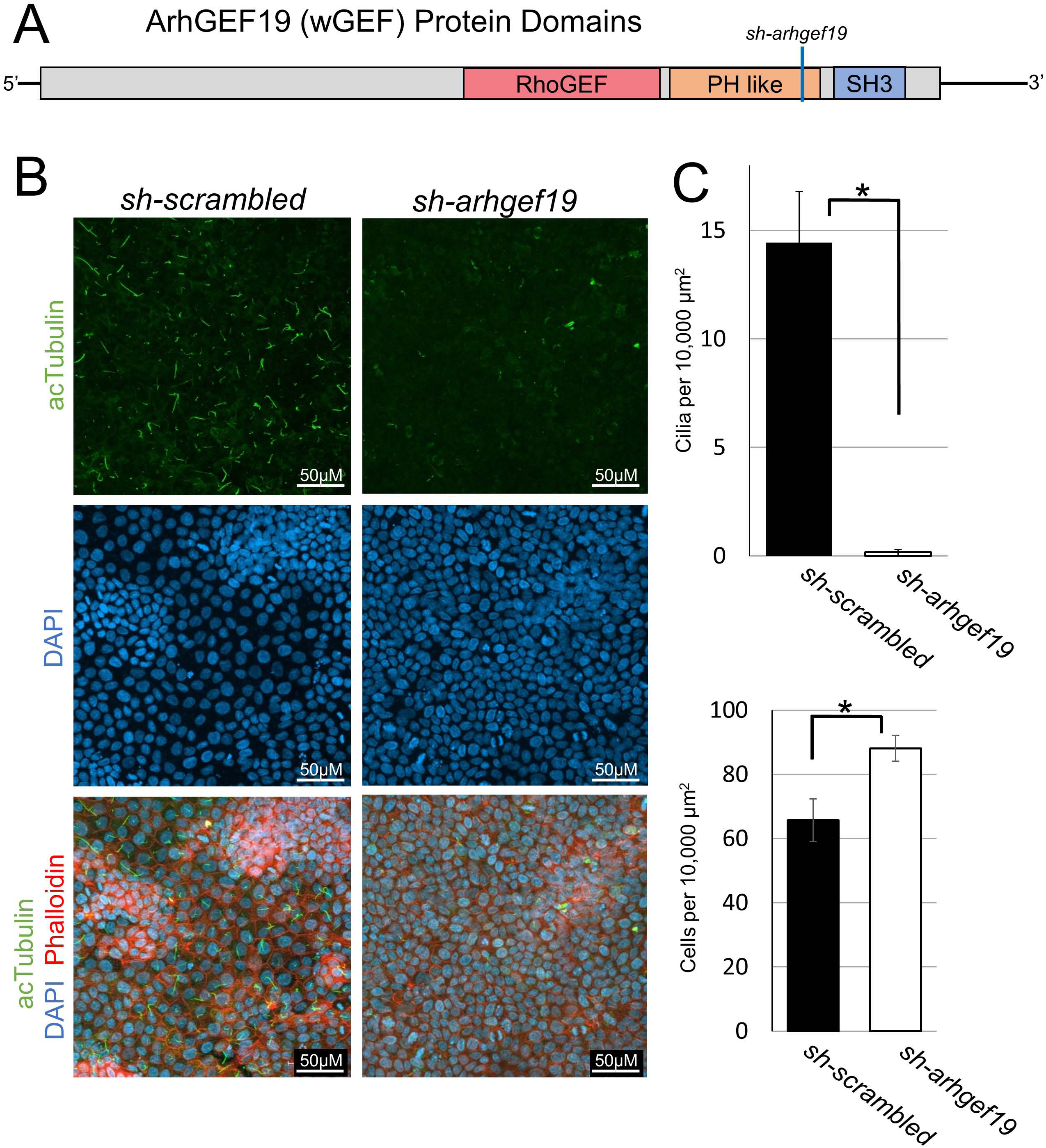
Arhgef19 knockdown results in loss of primary cilia in MDCKII cells. **A**) Diagram of the domains within the Arhgef19 (WGEF) protein. The corresponding position of the shRNA is marked with a blue line. **B)** MDCKII cells were infected with either a control construct or a construct that targets Arhgef19 and then polarized on transwell filters. Cells were stained with acetylated α-Tubulin antibody (acTubulin) to visualize primary cilia (green), DAPI to label nuclei (blue), and phalloidin to label F-actin (red). Confocal imaging was used to analyze the effects Arhgef19 depletion upon primary ciliogenesis. Scale bars equal to 50 μm. **C)** The number of cilia was measured, and cell numbers were quantified as described in Figure S5. * indicates p<0.05 as compared to *sh-scrambled*. Error bars are shown as ± SEM.

### Loss of Daam1 in *X. laevis* kidney does not cause loss of primary cilia

Daam1 activity has been shown to be required for proper nephron development in both *Xenopus* and zebrafish embryos (Miller et al 2011). Given that changes in nephron morphology and ciliogenesis are closely linked, *Xenopus* embryos were used to assess the role of Daam1 in formation of renal primary cilia. To avoid gastrulation defects associated with manipulating Daam1 expression in the whole embryo, a well-established kidney-targeted-morpholino approach was employed to knock down Daam1 activity in *Xenopus* nephric progenitors (Miller *et al.*, 2011; DeLay *et al.*, 2016; Corkins *et al.*, 2018). This approach consists of co-injecting Daam1 morpholino, or control morpholino, with mRNA encoding membrane-bound fluorescent protein (tracer to confirm correct delivery). Injections are made into selected blastomeres at the 8-cell stage that are fated to give rise to the kidney. As expected, Daam1 morpholino injected embryos show defects in nephron formation. Intriguingly, loss of primary renal cilia upon Daam1 knockdown was not observed (Figure 6), as was observed in MDCKII and IMCD3 cells following Daam1 depletion. While pronephric depletion of Daam1 leads to reduced elaboration of proximal and distal nephric tubules in stage 40 embryos, this phenotype is much less developed during early stages of nephron formation. To exclude the possibility that an earlier loss of renal tubules masks the loss of cilia phenotype, we also examined cilia in stage 30 embryos upon Daam1 knockdown. However, even in early stage embryos, there was no apparent loss of primary cilia upon Daam1 depletion (Figure S4). These data suggest that there are compensatory mechanisms that function *in vivo* to ensure proper ciliogenesis upon Daam1 knockdown.

**Figure 6:**
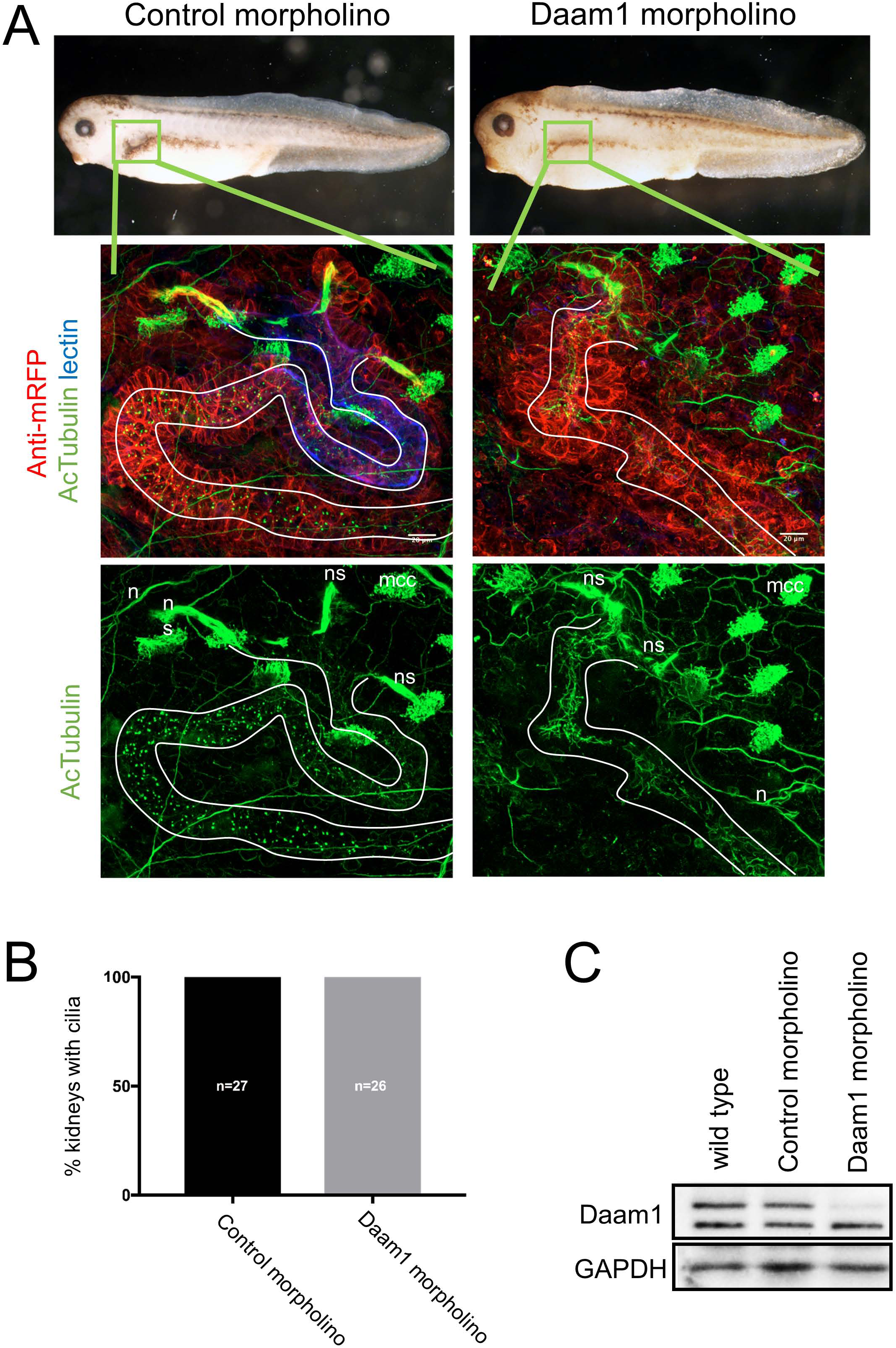
Daam1-depletion does not cause absence of cilia within *Xenopus* embryonic kidneys. Because knockdown of Daam1 in *Xenopus* kidney leads to kidney defects, we analyzed the effect of Daam1 knockdown on renal ciliogenesis. We injected Daam1 or Standard (Control) morpholino in combination with a membrane-tagged red fluorescent protein (mRFP) mRNA as a lineage tracer, into a *Xenopus* blastomere fated to the nephric anlagen. Embryos were fixed at stage 39-40 and stained with an antibody against acetylated α-Tubulin (acTubulin) to label cilia (green), anti-mRFP lineage tracer (red) and lectin to label theproximal tubule (blue). **(A)** Stereoscope brightfield imaging shows the gross morphology of Control and Daam1-morpholino injected embryos. Confocal fluorescent imaging of boxed regions (green) shows magnified views of corresponding kidneys. Kidney tubules displaying primary cilia are outlined in white. Neurons (n), multiciliated epidermal cells (mcc) and multiciliated cells within nephrostomes (ns) are immunostained with acetylated α-Tubulin (acTubulin) antibody. Scale bars equal to 20 μm. **(B)** The graph represents the percentage of Control (n=27) and Daam1-depleted (n=26) kidneys with primary cilia. **(C)** Western blot showing Daam1 protein expression levels in wild-type, control and Daam1 morphant embryos.

## DISCUSSION

Prior work demonstrated that Daam1 is important for the proper development of motile multiciliated cells (Yasunaga *et al.*, 2015). Daam1 forms the actin network for proper anchoring of the basal bodies. However, a role for Daam1 in formation of primary sensory cilia has not previously been identified. In this study, we analyzed the role of Daam1 in renal primary ciliogenesis.

Within the three models examined, the degree to which Daam1 influences ciliogenesis is variable. 2D *sh-daam1*-depleted MDCK cultures show absence of cilia, while 3D cultures have a weaker phenotype and display reductions in luminal ciliogenesis. *X. laevis* Daam1-morpholino depleted embryonic kidneys have renal cilia. There have been prior studies pointing to different observations arising as a function of the experimental model being employed (O’Brien *et al.*, 2001; Baek *et al.*, 2016). This dependence of phenotype upon methodology suggests that other factors may compensate for Daam1 and are likely to be influenced by extrinsic factors. For example, other formin proteins may facilitate actin polymerization to stabilize cilia in the absence of Daam1 *in vivo*. Alternatively, elements of the extracellular environment, such as extracellular matrix components, may compensate for Daam1 loss in embryos. Alternatively, given that Daam1 influences the positioning of motile cilia in *X. laevis* epidermal cells (Yasunaga *et al.*, 2015), it is possible that similar defects occur in renal primary cilia that are not be detectable by immunostaining.

It has previously been reported that Daam1 localizes to endocytic vesicles, actin filaments and the cell membrane (Kida *et al.*, 2007; Luo *et al.*, 2016). Based upon our finding that Daam1 localizes in vesicles that carry ciliary components, there are several potential mechanisms to explain how Daam1 plays a role in ciliogenesis. For example, Daam1 may be involved in vesicle trafficking to the base of the cilium or the docking of vesicles carrying ciliary components to the basal body. Alternatively, Daam1 be trafficked in vesicles to facilitate actin polymerization to stabilize ciliary structure at the base. Although we were not able to detect Daam1 at the primary cilia, it may only be transiently present or present at levels too low for us to detect. Daam1 is also necessary for proper centrosome orientation during cell migration. Ciliary basal bodies are modified centrosomes, so it is therefore possible that Daam1 performs an analogous function, in this case directing the orientation of the centrosome during ciliogenesis (Ang *et al.*, 2010).

The requirement of both ArhGEF19 and Daam1’s formin activity for ciliogenesis indicates a dual role for Daam1 in ciliogenesis through both actin polymerization and RhoA activation (Moseley *et al.*, 2006; Lu *et al.*, 2007). While future study will be required, it is possible that ArhGEF19 is functioning through a Daam1-independent mechanism to promote ciliogenesis. Given that no other proteins that affect ArhG19 activity have been identified and the effector proteins of Wnt-activated RhoA signaling are currently unknown, it is difficult to disprove this hypothesis. Our data indicate a similar cilia phenotype upon *daam1* and *arhgef19* depletion. However, *sh-daam1* cells tend to grow on top of each other, while *sh-arhgef19* cells do not, suggesting unique deficits in cell migration and epithelization in the *sh-daam1* cells. Thus, subsequent study will be needed to elucidate the role of ArhGEF19 in ciliogenesis.

Overall, our work indicates that the planar cell polarity effector, Daam1, promotes the formation of primary cilia in MDCKII and IMCD3 kidney epithelial cells. The formin activity of Daam1 and the Daam1-associated GEF, ArhGEF19, are required for proper cilia formation. Given prior work in *X. laevis* skin indicating Daam1 affects the polarized orientation of motile cilia (Yasunaga *et al.*, 2015), it is possible that in the kidney it is also affecting organization of the primary cilia. However, our data showing that Daam1 localizes in vesicles that carry ciliary components indicates a novel mechanism by which Daam1 may play a role in ciliogenesis.

## MATERIALS AND METHODS

### Cell culture

All cells were grown at 37°C with 5% CO2. Cells grown for experiments were passaged prior to confluency as confluent cells required excessive trypsinizing resulting in problems with ciliation. MDCKII cells were grown in DMEM + 10%FCS + Antibiotic/Antimycotic solution (Sigma A5955)(Williams *et al.*, 2017) and IMCD3 cells were grown in DMEM/F12 + 10%FCS + Antibiotic/Antimycotic solution (Mick *et al.*, 2015).

### Transfections and stable cell line generation

pLKO.1 lentiviral construct along with packaging vector ΔVPR and VSVG plasmids were transfected using Fugene-6 (Roche) or PEI (POLYSCIENCES 23966-2) into HEK293T cells grown on a 10cm dish (Moffat *et al.*, 2006). Supernatant containing transfection reagent was discarded and replaced with 5ml of fresh media. Supernatant was collected and replaced with 5ml fresh media over the course of 3 days resulting in 25ml of virus. The virus was filtered through 0.2μm filter to remove debris and frozen in 5ml aliquots at -80°C

Cells to be infected were split and allowed to grow to 50% confluency in a 10cm plate then washed with PBS. Virus was thawed at 37°C and 5μl 10mg/ml polybrene (Sigma H9268) was added to the virus, then the entire 5mls were added to cells. Cells were allowed to incubate with virus for 24 hours. Virus was then washed off and replaced with DMEM and the cells were allowed to grow for an additional 24hrs. DMEM +10μg/ml puromycin was then added to the cells and they were allowed to grow to confluency. Media was changed daily to remove cell debris.

Transient transfection was performed on uncoated glass coverslips or 6 well dishes using 5-μg of plasmid along with 30μg polyethylenimine, linear mw 25,000 (Polysciences 23966-2). Cells were washed with DMEM or DMEM-F12 2-4 hours after transfection reagent was added (Longo *et al.*, 2013).

### Ciliation

For MDCKII 2D cell culture, cells were diluted 1:10 from a nearly confluent plate onto plastic dishes and then allowed to grow overnight. 2.5×10^4^ cells were then plated onto transwell filters (Corning 3460) with DMEM on both sides of the filter. Cells were confluent the day after plating and were allowed to grow for an additional 4-5 days to ciliate. Media was changed every other day. For rescue experiments, cells were transfected in 6 well dishes and expression was verified prior to moving to transwell filters.

MDCKII cyst formation was modified from previous work (Elia and Lippincott-Schwartz, 2009; Hebert *et al.*, 2012). Cells were diluted 1:10 from a nearly confluent plate onto plastic dishes and then allowed to grow overnight. 8 well cell chambers (Falcon 08-774-26) were precoated with 85µl collagen solution [24mM glutamine, 2.35 mg/ml NaHCO_3_, 20mM HEPES (pH 7.6), 1x MEM, 2mg/ml type I collagen (Corning 354249)]. Cells were collected and strained through a 0.4µm strainer to remove cell clumps. 2.6×10^3^ cells were resuspended in 175μL collagen solution and placed in the precoated cell chambers. 400μL DMEM was add to each chamber after the collagen solidified. DMEM was changed once every other day for 12-14 days.

For IMCD3 cells, 2.5×10^4^ cells were plated onto glass coverslips and allowed to grow to confluency. Media was washed off and replaced with DMEM-F12 without serum for two days.

Cilia counts were performed by manually counting all cilia within an image. *sh-daam1* cells tend to pile on top of each causing automated methods of counting cells to miscount cells. For this reason, we split images up into a 4×4 grid and manually counted the same four sections in each image (Figure S5).

### Staining and imaging

#### 2D MDCK and IMD3 cell cultures

Cells were fixed in 4% PFA in PBS followed by 100mM Glycine. Cells were blocked in 10% goat serum in PTW (PBS+0.1% TritonX). 1:1000 Phalloidin-Alexa568 (Invitrogen, A12380) was used for staining of F-actin, and 1:1000 4’6’-diamidino-2-phenylindole were used to detect nuclei. Primary antibody 1:100 α-Tub1a (Sigma-T6793) in combination with secondary antibody anti-mouse IgG Alexa 647 (Invitrogen, A-21235).

#### 3D MDCK cultures

Staining of MDCK 3D cell cultures was carried out as previously described (Williams et al. 2017). Slides were incubated with primary 1:1000 mouse α-Tub1a (Sigma -T6793) antibody and detected with 1:200 anti-rabbit CY3 (Jackson Immunoreserach, cat# 715-165-151) or 1:200 Alexa 488 Mouse (1:200, Jackson Immunoresearch, cat# 715-545-150) secondary antibodies. F-actin was labeled with Alexa Fluor 647 Phalloidin (1:200, Invitrogen, A22287) while nuclei were detected with 4’6’-diamidino-2-phenylindole (1:500, DAPI). Slides were mounted in Flouromount-G medium (Southern Biotech, cat# 0100-01) prior to imaging.

#### *Xenopus* embryos

Embryos were staged (Nieuwkoop and Faber, 1994), fixed in MEMFA (DeLay *et al.*, 2016) and immunostained using established protocols (Hemmati-Brivanlou and Melton, 1994)Primary antibody 1:100 mouse α-Tub1a (Sigma -T6793), 1:250 rabbit RFP (MBL International -PM005) and 1:250 rabbit Lhx1 (gift from Masanori Taira, (Taira *et al.*, 1992)) were used. Proximal tubules were stained using fluorescein-coupled Erythrina cristagalli lectin at 50 #g/ml (Vector Labs).

Secondary antibodies anti-mouse IgG Alexa 647 (Invitrogen, A-21235), anti-mouse IgG Alexa 488 (Invitrogen, A-11001) and anti-rabbit IgG Alexa 555 (Invitrogen, A-21428) were used at 1:500 concentration. Embryos were dehydrated in methanol and cleared in a benzyl benzoate/benzyl alcohol (2:1) solution for imaging.

### Imaging

Images were taken using an Olympus SZX16 fluorescent stereomicroscope and an upright Leica SP5, inverted Nikon A1, and an inverted Zeiss LSM800 laser scanning confocal microscopes with cell incubation chamber. Nikon-Elements, Zen blue, ImageJ (Fiji plugin), Adobe Photoshop and Microsoft PowerPoint were used for data analysis and image processing.

For live imaging cell were grown on coverslips and imaged in Attofluor™ Cell Chamber (Thermo A7816).

### Western blot

#### Cell lysates

Cells were trypsinized from plates, collected and washed twice in PBS prior to being resuspended in 2X Laemmli (Biorad) plus 100 µM dithiothreitol. The resulting cell lysates were then boiled at 95°C for 30min. Lysates were run on an 8% SDS-PAGE gel and the protein was then transblotted onto a 0.2 µm nitrocellulose membrane (GE Healthcare), followed by blocking for 3 hours in KPL block (SeraCare) at room temperature. Blots were incubated in 1:1000 rabbit anti-GAPDH (Santa Cruz sc-25778), 1:1000 rabbit anti-Daam1 (Protein Tech 14876-1-AP) or 1:1000 rabbit anti-β-catenin (McCrea *et al.*, 1993) primary antibodies for 1-2hrs. Blots were then washed with TBST, incubated in goat anti-rabbit IgG horseradish peroxidase secondary antibody (1:5000, BioRad) for 2 hours at room temperature and then washed again with TBST prior to imaging on a BioRad ChemiDoc XRS+ imaging system using enhanced chemiluminescence (Pierce Supersignal West Pico). Preparation of MDCK cell extracts and western blot analysis presented in Figure 2 were carried out following the protocol described by (Williams *et al.*, 2017b).

#### Embryo lysates

1-cell *Xenopus* embryos were injected with 10nl of 20ng Daam1 or Standard morpholino in combination with 0.5 ng mRFP. Lysates were prepared from Stage 11 embryos and western blots were performed following previously established protocol (Miller *et al.*, 2011).

### Plasmids

pCS2-GFP-Daam1 and pCS2-GFP-Daam1(I698A) constructs were a gift from the Goode lab(Moseley *et al.*, 2006; Lu *et al.*, 2007). A mutation of A2822G was discovered in these plasmids, which resulted in the amino acid change of D941G. This mutation was corrected using site directed mutagenesis. The pCS2-GFP plasmid was generated by digesting pCS2-GFP-Cby1 (A gift from the Klymkowski lab) with ClaI - XbaI to remove the *cby1* gene (Shi *et al.*, 2014). The ends were blunted using phusion DNA polymerase (NEB) followed by ligation and transformation.

pCS2-mCherry-Daam1 and pCS2-mCherry-Daam1(I698A) constructs were generated by BamHI-NotI digestion of pCS2-GFP-Daam1 and pCS2-GFP-Daam1(I698A) and cloned into the same sites of pCS2-GFPCby1 replacing the GFPCby1 with Daam1. mCherry was PCR amplified from Cas9-mCherry (Addgene #78313) and cloned into the BamHI site. *sh-daam1* (#1 TTT<u>C</u>AGGAGA<u>T</u>AGTATTGTGC, #2 TAACATCAGAAATTCATAGCG, #3 AAACAGGTCTTTAGCTTCTGC) and *sh-arhgef19* (TGCTTCTCACTTTCGGTCC) constructs were purchased from GE-Dharmacon. *sh-scrambled* plasmid was used as a negative control for 2D ciliation experiments. As *sh-daam1* (#1) construct is designed against the human *daam1,* and perfectly matches the *Canis lupis daam1* however, the target sequence is a two base pairs off of the mouse daam1 sequence. Thereforea *sh-daam1* (*ms #1*) (TTT<u>T</u>AGGAGA<u>A</u>AGTATTGTGC) construct was generated to better target *daam1* in IMCD3 cells. The construct was made by cloning oligos containing a hairpin for the indicated sequence into EcoRI - AgeI sites of pLKO.1 as described in (Moffat *et al.*, 2006).

#### Xenopus laevis

Wild type oocyte-positive *X. laevis* adults were purchased from Nasco (LM00531MX) and embryos were obtained from these adults and reared as previously described (DeLay *et al.*, 2018). This protocol was approved by the University of Texas McGovern Medical School Institutional Animal Care and Use Committee (IACUC) (protocol #: AWC-16-0111). Microinjections were performed as previously described (Sive *et al.*, 2000; DeLay *et al.*, 2016). 10 nL of injection mix was injected into the indicated blastomere. For morpholino injections, 20 ng of Daam1 morpholino (5-GCCGCAGGTCTGTCAGTTGCTTCTA 3-) (Miller *et al.*, 2011; Corkins *et al.*, 2018) or Standard morpholino (5-CCTCTTACCTCAGTTACAATTTATA 3-) were injected along with 500 pg of membrane targeted RFP RNA as a tracer to mark targeted cells into the V2 blastomere at the 8-cell stage to target the kidney (Moody, 1987).

## Acknowledgements

We appreciate the helpful suggestions and advice throughout this project from the members of the laboratories of R.K. Miller, A.B. Gladden, P.D. McCrea, as well as M. Kloc, especially to Bridget D. Delay critical evaluation of this manuscript. We thank the animal care technicians and veterinarians, including J.C. Whitney and T.H. Gomez who took care of the animals even during Hurricane Harvey. For imaging support, we thank the Department of Pediatrics imaging core supported by the Office of the President of Academic Affairs, as well as Adriana Paulucci and the MD Anderson Genetics Imaging Core. We thank all of the labs that provided plasmids for this project. pCS2-GFP-Cby1 was a gift from Mike Klymkowsky (University of Colorado Boulder), and pCS2-GFP-Daam1 and pCS2-GFP-Daam1(I698A) were a gifts from Raymond Habas (Temple University) and Bruce L. Goode (Brandeis University), respectively. pCS10R-IFT88 was a gift from John Wallingford. We also thank Masanori Taira for Lhx1 antibody.

These studies were supported by the National Kidney Foundation (FLB1628 to R.K.M), National Institutes of Health (NIH) grants (K01DK092320 and R03DK118771 to R.K.M.), and startup funding from the Department of Pediatrics, Pediatric Research Center at UTHealth McGovern Medical School (to R.K.M.).

## Authorship contributions

M.E.C, V.K.S., R.K.M, and A.B.G generated *sh-daam1* cells and performed 2D cilia experiments. A.B.G., V.K.S. and M.E.C. preformed 3D cyst experiments. V.K.S. and M.E.C. generated plasmids for viral infections, GFP-Daam1 and GFP-Daam1(I698A) plasmids for transient transfections and wrote the manuscript. M.E.C generated mCherry-Daam1 and mCherry-Daam1(I698A) plasmids for transient transfections and performed experiments to generate figures 1,3,4,5, S1, S2, and S5. V.K.S and R.K.M. performed experiments for *X. laevis* kidney experiments. V.K.S. generated data for figures 2, 6, S3, and S4. R.K.M., M.K. and V.K.S. performed experiments for Daam1 morpholino knockdown in *X. laevis* skin. P.D.M. facilitated the initiation of this project. A.B.G helped set up mammalian cell culture and all cell culture techniques, oversaw experiments performed in cell culture. R.K.M. conceived of the project and performed preliminary experiments. All authors have provided critical feedback in the writing of the manuscript.

---

**Figure S1:**
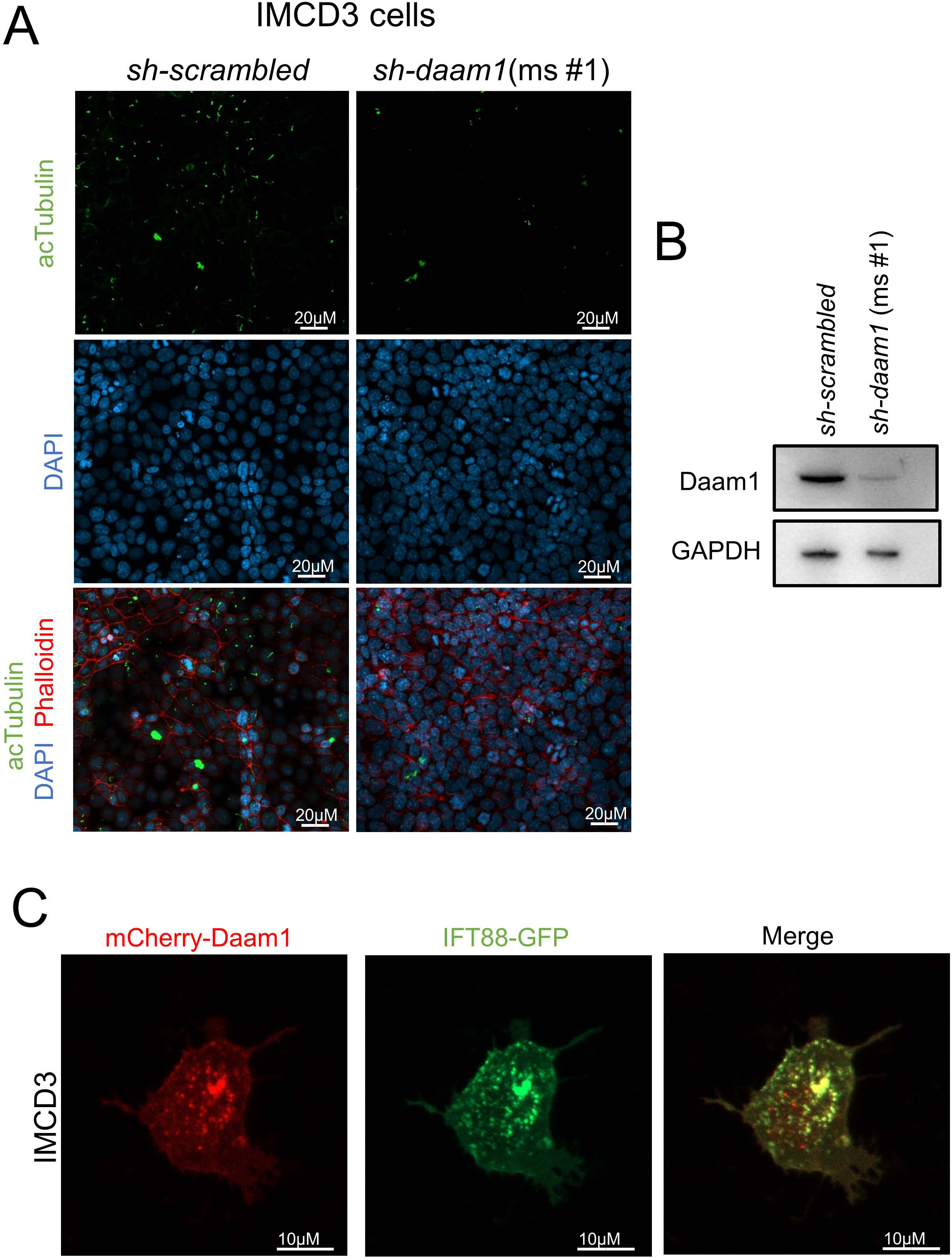
Loss of Daam1 results in a reduction of primary cilia and mCherry-Daam1 localizes to vesicles carrying IFT88 in IMCD3 cells. Murine inner medullary collecting duct (IMCD3) were infected with either *sh-daam1* or a control construct then ciliated on glass coverslips. **A)** Cells were stained with acetylated α-Tubulin antibody (acTubulin) to label primary cilia (green), DAPI to label nuclei (blue), and phalloidin to label F-actin (red). Confocal imaging was used to analyze the effects Daam1 depletion upon primary ciliogenesis. Scale bars equal to 20 μm. **B)** Western blot of *sh-daam1* IMCD3 cell lysates showing depletion of Daam1 protein levels. GAPDH was used as a loading control. **C)** Cells were co-transfected with constructs that express mCherry-Daam1 and IFT88-GFP than imaged in live cells. Scale bars equal to 10 μm.

**Figure S2:**
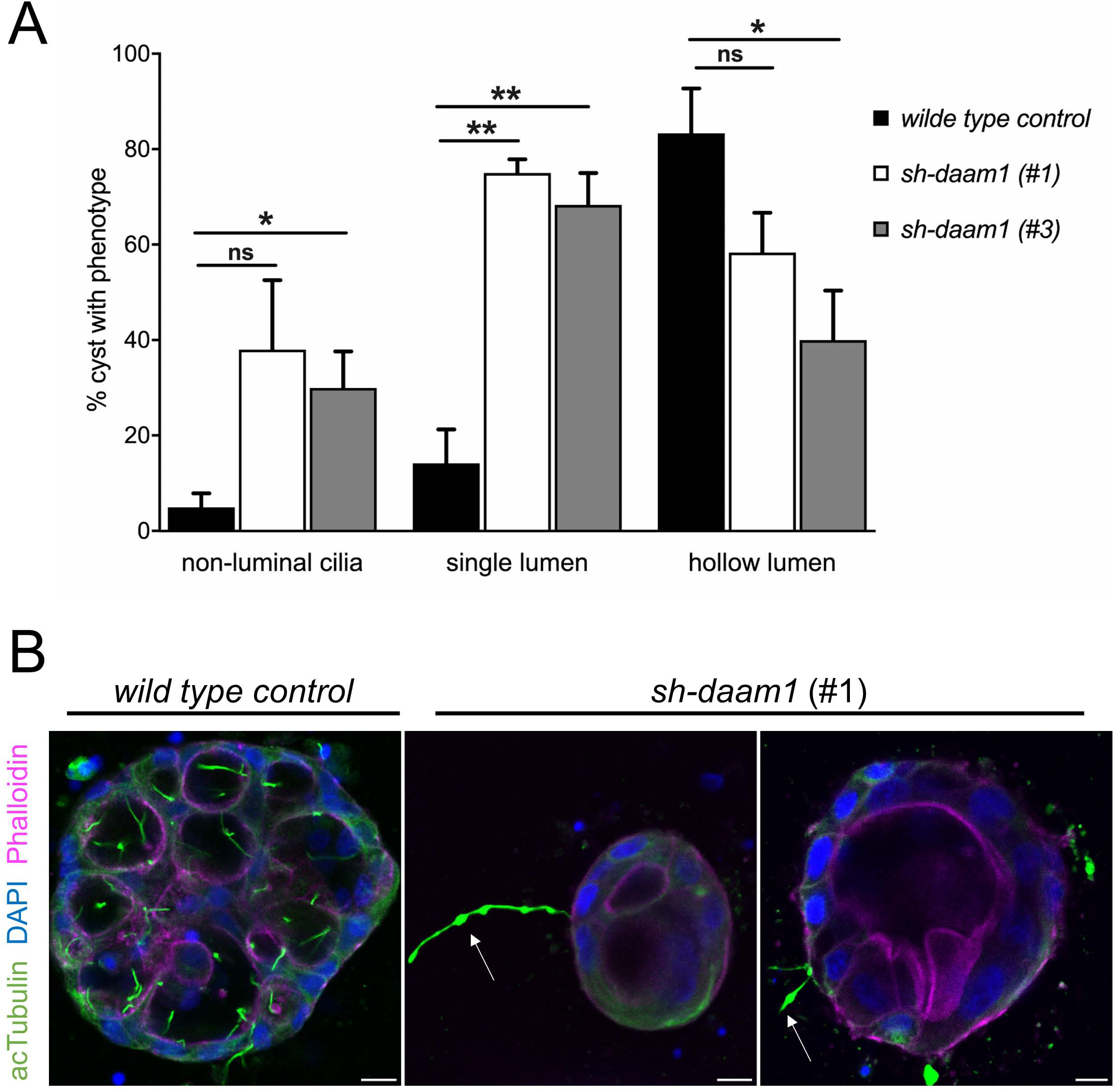
Phenotypes derived from control and *sh-daam1* knockdown. Daam1-depleted 3D MDCKII cyst were scored for the presence of (1) non-luminal cilia - cilia that do not protrude into central lumen, (2) multiple lumens and (3) hollow lumens-luminal clearance. Twenty cysts were randomly selected for analysis in three independent experiments. **(A)** The graph indicates the relative percentage of cyst for each phenotype. Error bars are shown as ± SEM; Significance was calculated using unpaired, two-tailed t-test; ns indicates p>0.05, * indicates p<0.05, **p<0.01 **(B)** Representative images of cysts with non-luminal cilia phenotype. In Daam1 depleted cysts, white arrows point at cilia protruding out into extracellular matrix. Scale bars equal to 10 μm.

**Figure S3:**
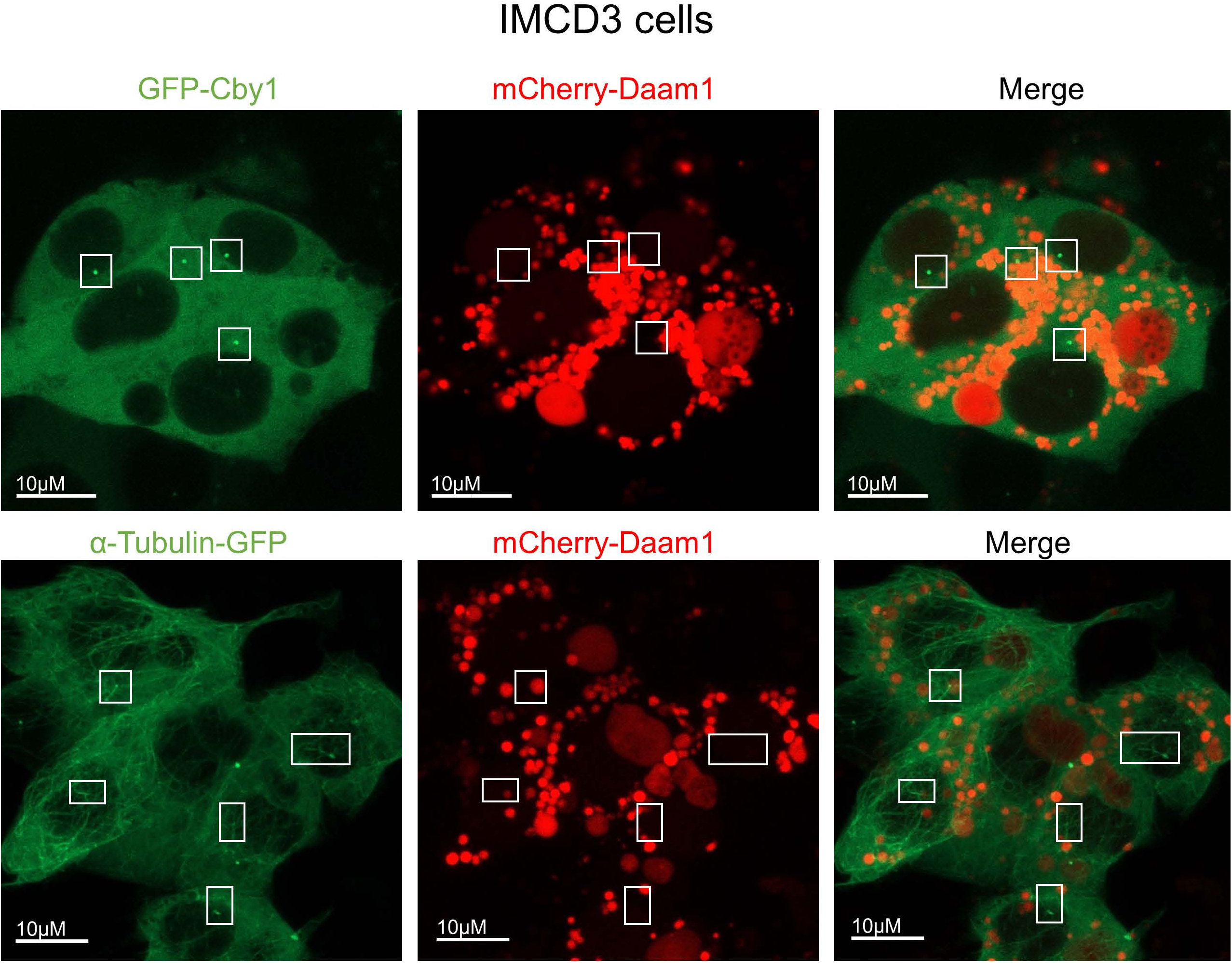
Daam1 does not localize to the primary cilia in IMCD3 cells. Murine inner medullary collecting duct (IMCD3) cells were transfected with mCherry-Daam1 along with either Cby1-GFP or α-Tubulin-GFP. Cells were grown to confluency and serum starved to ciliate. Then cells were analyzed via confocal for colocalization of Daam1 and these ciliary markers. White boxes outline transition zone in Cby images and cilia in α-Tubulin images. Scale bars equal to 10 μm.

**Figure S4:**
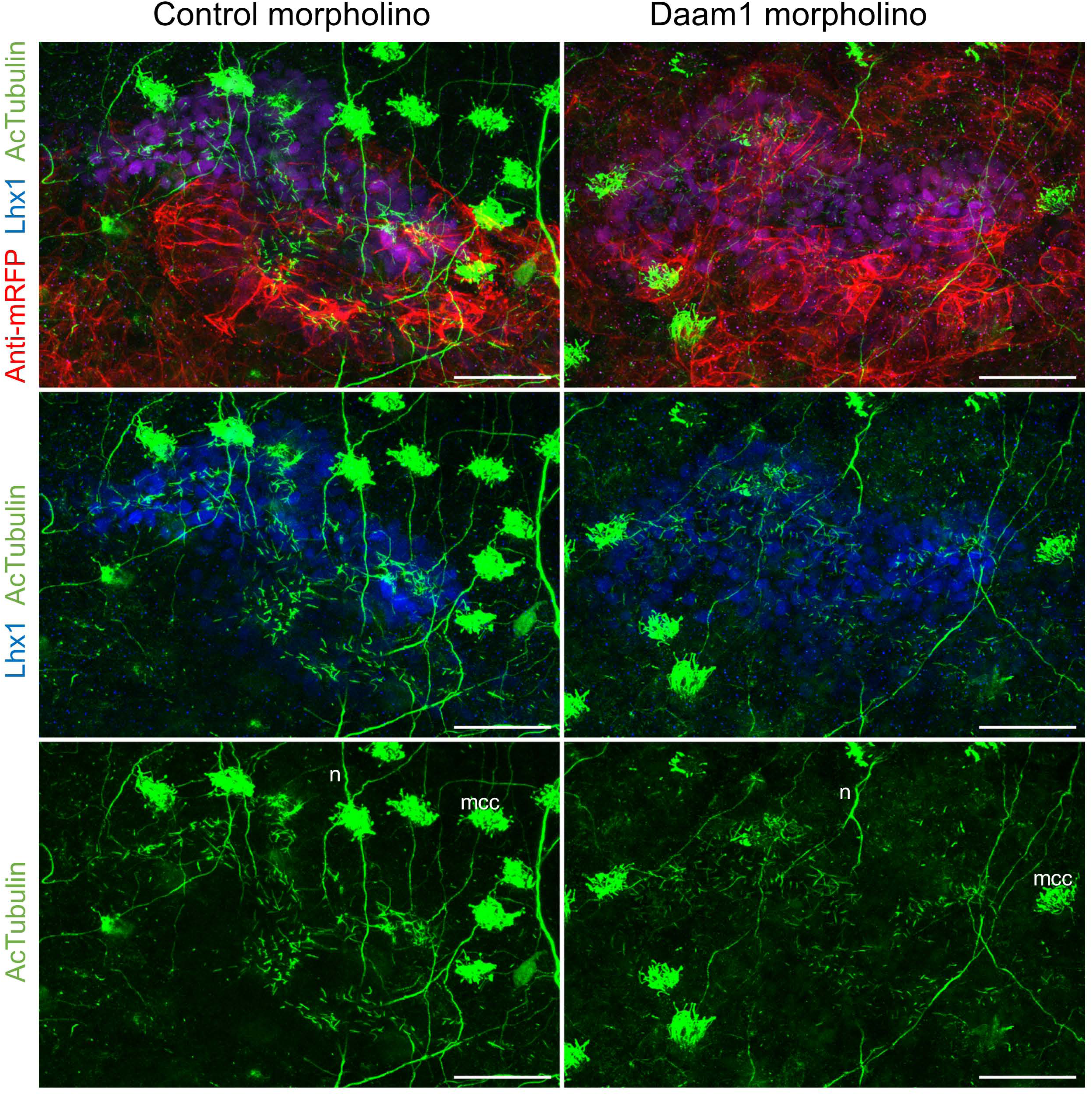
Daam1-depletion does not lead to the absence of cilia during development of *Xenopus* embryonic kidneys. To further analyze the effect of Daam1 depletion on ciliogenesis, we fixed 8-cell Daam1 and Standard (control) morpholino injected embryos during early stages of kidney morphogenesis (stage 30). mRFP mRNA was used as a lineage tracer and coinjected with morpholinos. Stage 30- fixed embryos were immunostained with an antibody against anti-mRFP to visualize tracer (red) together with an Lhx1 antibody to label nephric progenitor cells (blue) and acetylated α-Tubulin antibody to label primary cilia (green). Subsequently, embryos were analyzed using a confocal laser-scanning microscope and representative maximum projections of Z-stack sections are shown. Acetylated α-Tubulin antibody also stains neurons (n) and multiciliated epidermal cells (mcc). Scale bar is equal to 50 μm.

**Figure S5:**
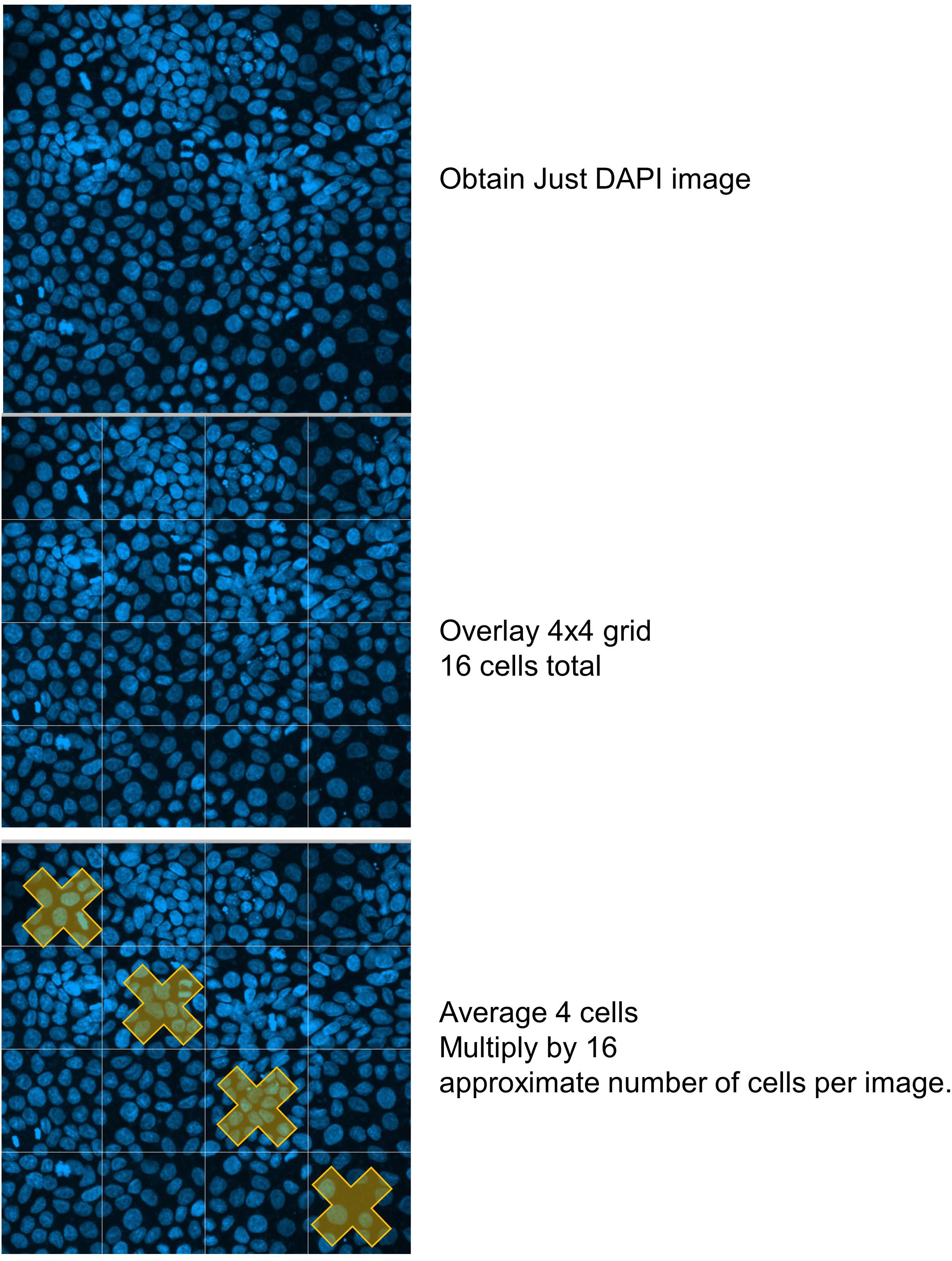
Diagram demonstrating the methodology used to count number of cells. To obtain an unbiased quantitation of cell numbers in MDCKII depletion experiments (Figures 1 and 5), DAPI images were divided into a 4×4 grid and cells were counted within the 4 cells along the diagonal from the upper left to the lower right of the image and the number of cells was averaged. This number was multiplied by 16 to obtain the approximate number of cells per image.

